# Coding Recognition of the Dose-Effect Interdependence of Small Biomolecules Encrypted on Paired Chromatographic-Based Microassay Arrays

**DOI:** 10.1101/2021.03.10.434720

**Authors:** Yifeng Deng, Zhenpeng Lin, Yuan Cheng

## Abstract

The discovery of small biomolecules suffers from the lack of a comprehensive framework to express the intrinsic correlation between bioactivity and contributing small molecules in complex samples with molecular and bioactivity diversity. Here, by mapping a sample’s 2D-HPTLC fingerprint to microplates, the paired chromatographic-based microassay arrays are created, which can be used as quasi-chip to characterize multiple attributes of chromatographic components; and as the array differential expression of the bioactivity and molecular attributes of irregular chromatographic spots for dose-effect interdependent encoding; and also as the automatic-collimated array mosaics of the multi-attributes of each component itself encrypted by its chromatographic fingerprint. Based on this homologous framework, we propose a correlating recognition strategy for small-biomolecules through their self-consistent chromatographic behavior characteristics. In the approach, the small-biomolecule recognition in diverse compounds is transformed into a constraint satisfaction problem, which is addressed through examining the dose-effect interdependence of the homologous 2D code pairs by array matching algorithm, instead of preparing diverse compound monomers of complex test sample for identifying item-by-item. Furtherly, considering the dose-effect interdependent 2D code pairs as links and the digital-specific quasimolecular ions as nodes, an extendable self-consistent framework that correlates mammalian cell phenotypic and target-based bioassays with small biomolecules is established. Therefore, the small molecule contributions and the correlations of bioactivities, as well as their pathway can be comprehensively revealed, so as to improve the reliability and efficiency of screening. This strategy was successfully applied to galangal, and practiced the high-throughput digital preliminary screening of small-biomolecules in a natural product.

**Highlight:** Rapid self-recognition of small biomolecules in diverse samples without pre-isolation Recognized by dose-effect interdependency developed from homologous TLC fingerprint Matching of HPTLC-based molecular imprinting and bioautography on microassay arrays Microarray-based differential expression of substance attributes instead of spot scan An array framework for combining phenotype-based and target-based assays with TLC-MS

**ToC graphic:** 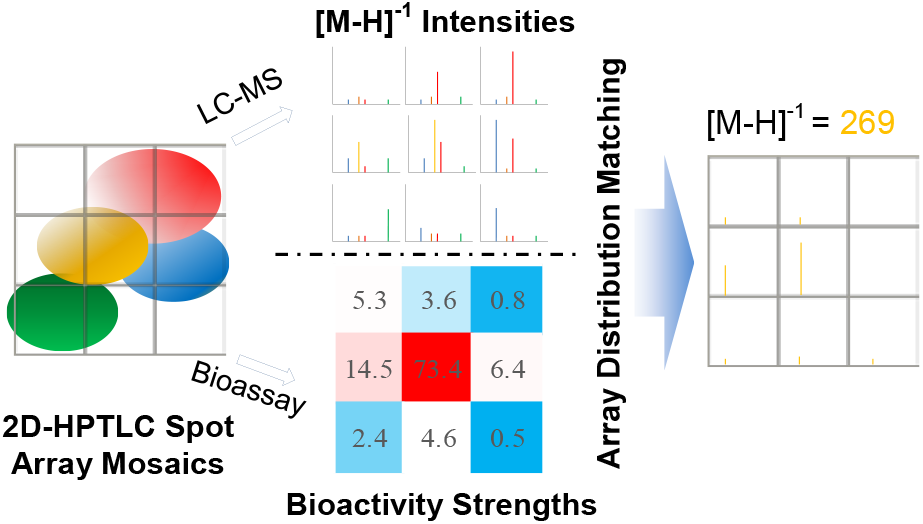

## Introduction

The identification of small molecules, including diverse compounds with novel chemical structures, diagnostic markers and drugs, plays an important role in drug discovery and precision medicine [1, 2]. As new coronavirus varieties become more and more rampant, people have great hopes for early discovery of small molecule drugs against coronaviruses. Although in recent years, advances in omics technologies have begun to enable personalized medicine at an extraordinarily detailed molecular level [3, 4], while research on small biomolecules seems to be much slower. Since phenotype-based assay and target-based assay and chemical identification were performed respectively, drug development is plagued by technical uncertainty and associated risks of failure [5, 6]. How to “integrate both the phenotypic and target-based approaches to estimate a relevant network from compound to phenotype in screening” becomes an emerging challenge [7, 8].

An earlier well-known bio-directed assay for small molecules in diverse compounds was bioautography on thin-layer chromatography (TLC) [9, 10]. Recently, Morlock et al. immersed the HPTLC (high-performance thin-layer chromatography) plates with the separated sample extracts in the bacterial culture medium and extracted the components in the bioactive spots through a special TLC-MS interface (Camag) for *in situ* biotracing [11]. However, due to the interference of the chromatographic matrix on many biological assays, the application of this method has been limited and has not been applied to mammalian cells. Meanwhile, electrophoresis, another type of planar chromatography, has become an important means of omics, because it can be combined with macromolecular immune reactions by transferring gel electrophoresis strips to the electrophilic membranes. Molecular imprinting derived from chromatography, molecular recognition based on binding the template to the functional monomer, is widely used in the fields of chromatographic separation, but it is limited to unidirectional matching of a preset specific target molecule [12]. In recent decades, crafted molecular diversity libraries and various cutting-edge microchips, such as small molecular microarrays and microfluidic chips [13, 14], have emerged; they are being used for various bioassays or chemical analysis with high throughput and easy automation features [15]. However, these assays are highly dependent on molecular diversity libraries and are seldom used for complex samples, such as natural products and clinical samples. Since it is difficult to capture the diversity of molecules of real samples, and the diverse compounds of a complex samples have to be separated to monomers regardless of whether they are bioactive [16]. On the other hand, mass spectrometry imaging (MSI) has emerged as the future frontiers of chemical analysis [17], which combines the molecular and structure information gained from mass spectrometry with visualization of spatial distributions in bio-sample sections. Now, thin layer chromatography coupled with mass spectrometry (TLC–MS) has developed into one of the most efficient analytical tools for chemical identification and structural elucidation of bioactive analytes on TLC [18, 19]. As such, establishing a comprehensive framework to express the “intrinsic correlation” between bioactivity and contributing small molecules in complex samples with molecular and bioactivity diversity remains a prominent issue [7, 8].

With the success of today’s omics research, correlation recognition methods for biomacromolecules in complex samples play an important part. The most important aspect of these methods is the specific correlation between molecular characteristics and bioactivities, such as the pairing of bases encoded on DNA arrays and antigen-antibody pairing based on the “molecular recognition exhibits molecular complementarity” practice [20, 21]. However, correlation recognition methods in nature are far beyond these. Macroscopically, NASA’s night-light image pair of the Earth has been used to map the human socioeconomic activity [22, 23]. Inspired by these approaches of correlating specific behavioral observation, we realize that the homologous chromatographic behavior characteristics of a small molecule can also be used to recognize multiple attributes derived from itself, such as molecular characteristics and modulating bioactivity, and thus propose a correlating recognition strategy for small molecules with bioactivity based on homologous 2D-HPTLC fingerprint.

Around this research strategy, this article discusses the following issues: characterizing multiple attributes of chromatographic components in irregular chromatographic spots as microarray distribution gradients, transforming the microassay array results into two-dimensional codes and self-recognizing biomolecules with array matching algorithm, etc., and the scientific validity, the error control and the affecting factors of this strategy are also discussed.

## Material and methods

### Materials and reagents

Galangal (Alpinia officinarum Hance) was Collected in Longtang Town, Guangdong Province, China. Galangin and Diphenylheptane A was purchased from NICPBP (Lot No. 111699-2006021 and 11757-200601). 5-Fluorouracil was purchased from Shanghai EKEAR (Lot No. 150729). Gold Nanoparticles (GNPs) prepared with reference to the literature [24]. AlamarBlue™ Cell Viability Reagent was purchased from Invitrogen (Cat#DAL1025). Human A549 cells and Human HepG2 cells was provided by Clinical Research Center of Guangdong Medical University. Oligonucleotides (G-DNA): 5’-TTA GGG TTA GGG TTA GGG TTA GGG-3’ was synthesized by Sangon Biotech (Shanghai Bioengineering Co., Ltd., order No. 300064096). The reagents used in the experiment were all analytical reagents.

### Establishment of a 2D-HPTLC fingerprint developed in the microarray format

Dry powder (250 μm) of *Alpinia officinarum* Hance was extracted by conventional ultrasonication with petroleum ether, ethanol, tetrahydrofuran, acetone, dichloromethane, ethyl acetate, benzene and water (each representing one of the 8 kinds of solvents classified by Snyder). The extracts were combined to obtain a representative extract of galangal.

The sampling amounts were chosen by conventional pre-bioassays of the crude sample extract. Generally, the spotting volume of the crude extract on the HPTLC plate is controlled to be no more than 10 μL. The selected amount of the extract was spotted to achieve a 2.5 mm diameter on an HPTLC plate. HPTLC was extended to 7.2 cm × 7.2 cm according to the 384-well microplate array format (4.5 mm×4.5 mm matrix; the opening of the well of a 384-square-well microplate is approximately the size of a typical HPTLC spot), and the chromatographic two-dimensional mobile phase was used to establish a 2D-HPTLC fingerprint. The optimized two-dimensional mobile phase [25] used for 2D-HPTLC consisted of trichloromethane/MeOH/petroleum ether at a ratio of 7.76/0.24/2 (v/v/v) and ethyl acetate/petroleum ether/acetic acid at a ratio of 3/7/0.2 (v/v/v).

### Chromatographic-based microassay array preparation

According to the microarray format of the high-throughput microplate, the silica gel layer of the 2D-HPTLC fingerprint is defined and differentiated by the microarray into a square chip array [26], and the components on the square chip array are mapped into a microplate as a chromatographic-based microassay array with chromatographic matrix removed. Specifically, the silica gel layer was split and stripped from a thin aluminum sheet with a metal grid interface having a 384-well microplate array format (3D printing, original equipment manufacturer (OEM) according to the drawing, Supporting Information S1) under a parallel pressure of 8×10^3^ kPa (Specac Atlas® Manual Hydraulic Press-15T) to fabricate a flat square chip array (Figure 1, Supporting Information S2). The chip array was kept in the array arrangement as a whole, positioned in the corresponding 384-well filter plate, and then the components were positioned and eluted into the corresponding well of another 384-well microplate with methanol. Note that the G-quadruplex ligand bioassay should be pre-washed three times with 15 μL of water per well to remove possible salts. The microplate containing the eluent was evaporated to dryness in a vacuum centrifuge concentrator to prepare a stock chromatographic-based microassay array (Supporting Information S3). A schematic diagram of the preparation and workflow of the chromatographic-based microassay arrays is presented (Figure S3). The paired chromatographic-based microassay array with the consistent chromatographic distribution is prepared by transferring a portion of the sample from the stock chromatographic-based microassay arrays to the corresponding array units of another 384-well microplate.

**Figure 1.**
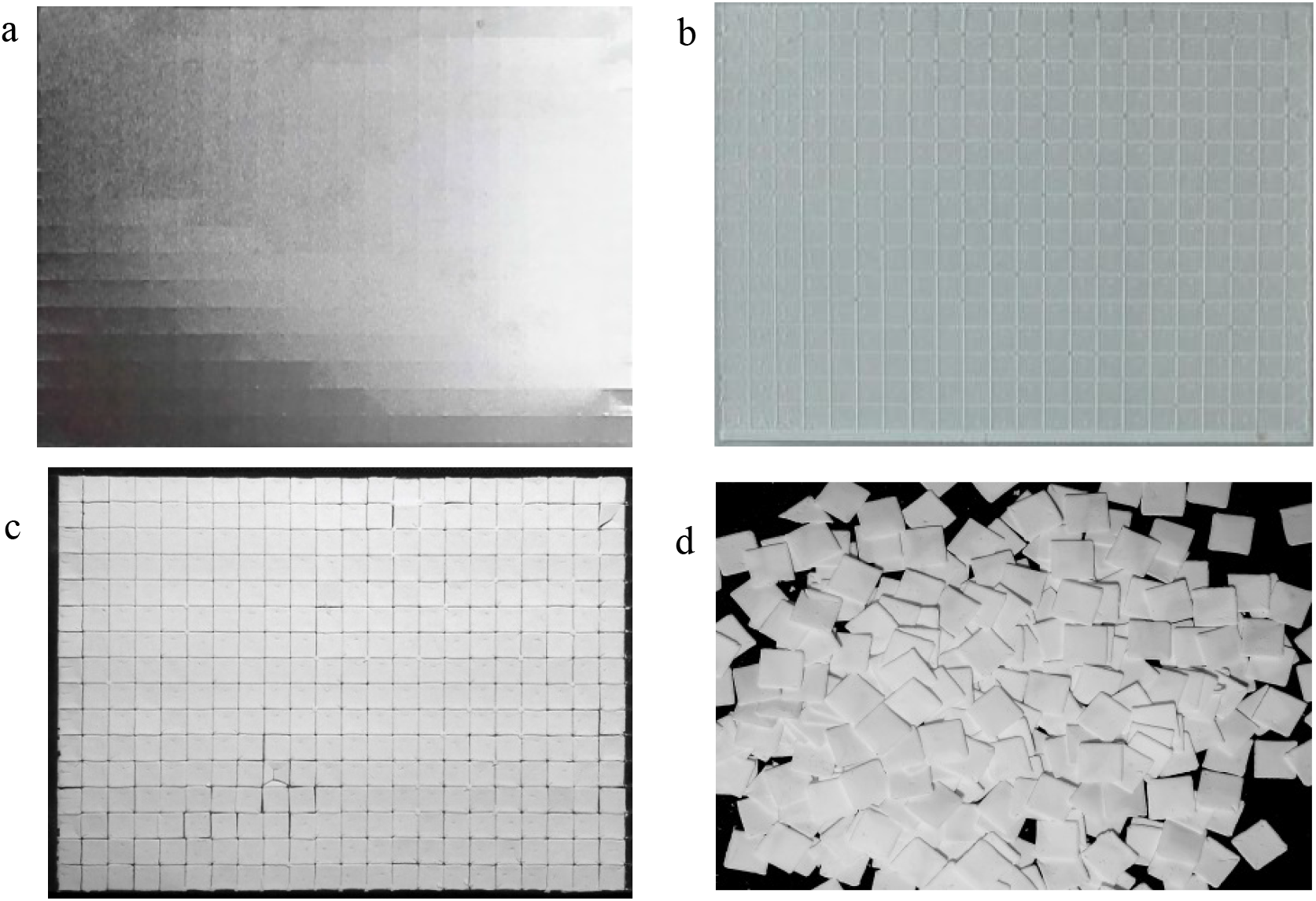
Array division and stripping effect of a HPTLC fingerprint. (a) Front and (b) back sides of the aluminum HPTLC sheet that had been pressed by the metal grid interface, (c) square silicon layer array stripped off and adhered to a PVDF film (painted black), and (d) scattered square silica gel layer chips.

### Cell proliferation assays

The proliferation assay of human A549 and HepG2 tumor cell lines is adopted for screening anticancer drugs [27]. The A549 and HepG2 cell lines were obtained from the Affiliated Hospital of Guangdong Medical University. With reference to Invitrogen’s AlamarBlue® Cell Viability Assay Protocol and related literature methods [28, 29], three groups of experiments were set up as follows: blank group, positive control group and sample group. Then, 0.1% DMSO and 1% glycerol were used as the reagent blank group, while 5-fluorouracil at a concentration of 0.077 mmol•L^-1^ was used as a positive control. The amount that adequately reflected the bioactivity of the chromatographic-based microassay array sample was chosen through the cell viability pre-bioassay as the test dose for subsequent bioassays [25].

The cells were cultured in high-glucose DMEM with 8% fetal bovine serum at a cell density of 1,000 cells per 40 μL in each well of the 384-well microplate. The cells were incubated at 37°C under 5% CO_2_. At 12 hours, 20 μL of the cell culture medium containing the optimal amount of test sample was added to the corresponding well of the microplate, and the cells were further incubated for 36 hours. Then, each group was centrifuged (12 ×g, 15 s) to remove the drug-containing cell culture medium and the cells were washed twice with PBS and discarded after centrifugation. Fifty microliters of cell culture medium and 5 μL of AlamarBlue were added per well to each group, and the samples were degassed with a vacuum centrifugal concentrator (269 ×g, room temperature, -80 KPa, 1 min). The absorbance of the microplate was measured at 570 nm and 600 nm (Bio-Tek Epoch one). Then, the cells were further incubated at 37°C under 5% CO_2_ for 12 hours. The absorbance was measured again using the same method. The difference between the absorbance of the cells incubated 12 hours after the addition of AlamarBlue and the absorbance at the time of initial addition was used as the absorbance value, and the cell survival rate was calculated according to Invitrogen’s protocol for the AlamarBlue® Cell Viability Assay.

### G-quadruplex ligand bioassay

G-quadriplex ligand bioassay is adopted for screening anticancer drugs [30]. Referring to the literature methods and with slight changes [31, 32], we used gold nanoparticles (GNPs) as a colorimetric probe and screened G-quadruplex ligands by colorimetry based on the color differences between single-stranded DNA-GNPs and G-quadruplex-GNPs. The amount that adequately reflected the bioactivity of the chromatographic-based microassay array sample was chosen through the G-quadruplex ligand pre-bioassay as the test dose for subsequent bioassays. The initial total volume is 35μL per well, which contains 5μL methanol. As previously described in the literature [24], the GNPs were prepared by reducing chloroauric acid with sodium citrate. The diameter was calculated to be 12 nm, and the concentration was 9.45 nM. The GNPs and 1.0 μM GDNA (5’-TTA GGG TTA GGG TTA GGG TTA GGG-3’ were mixed at a 1:150 molar ratio (optimized through an anti-NaCl-induced aggregation experiment performed by microplate titration, in which the range of the molar ratio of GDNA to GNPs was from 50-250) and incubated at room temperature (25°C) for 16 hours to prepare the GNP-GDNA probe. Then, 60 μL of the GNP-GDNA probe per well was added to the chromatographic-based microassay array. After 3 hours of incubation at room temperature (25°C), NaCl solution was added to each well in the microplate containing the reaction solution to achieve a concentration of 0.106 M, which was followed by further incubation for 30 min at room temperature (25°C). After the solution was added, the microplate was treated with a vacuum centrifugal concentrator (269 × g, at room temperature, -80 kPa, 1 min) to remove bubbles. The absorbance spectra (400∼850 nm) of each well of the microplate were collected (Bio-Tek Epoch one). The color change was observed, and the absorbance ratio calculated at 670 nm and 520 nm was used to evaluate the ligand activity of the components in each array unit of the chromatographic-based microassay array.

### LC–ESI-MS analysis

According to the bioactivity heat map, a portion of the sample corresponding to the significant bioactive array units is transferred from the stock chromatographic-based microassay array to the corresponding array units of another 384-well microplate for LC-ESI-MS analysis, which was conducted with an Agilent 6430 triple quadrupole mass spectrometer coupled with an electrospray ionization (ESI) source. The operating conditions were as follows: gas flow rate of 12 L•min^-1^, gas temperature of 350°C; sheath gas flow rate of 12 L•min-1, sheath gas temperature of 350°C and capillary voltage 3.6 kV. The mass analyses were performed with an ESI source in negative ionization mode; the m/z scan range was set from 65 to 750. The chromatographic system was composed of an Agilent 1200 series HPLC. The eluent was 75% methanol-H_2_O introduced at a flow rate of 0.2 mL•min^-1^. The injection volume was 5 μL in direct injection mode. Agilent Mass Hunter software was used for data analysis.

## Results and discussion

### Protocol design for the correlation between molecules and bioactivities

The paired chromatographic-based microassay arrays are created from an array-differential sample’s 2D-HPTLC fingerprint with the chromatographic matrix removed and the consistent chromatographic distribution (Supporting Information). This paired chromatographic-based microassay arrays can be used as quasi-chip to characterize multiple attributes of chromatographic components [33], and as the array differential expression of the bioactivity and molecular attributes of irregular chromatographic spots for coding interactions [26], as well as the automatic-collimated array mosaics of the multi-attributes of each component itself encrypted by its chromatographic fingerprint. A bioassay, i.e., a cell-based phenotypic assay or ligand binding assay, is performed using one of the paired chromatographic-based microassay arrays, and biotracing is followed correspondingly by direct ESI-MS analysis on the other one. Thus, local self-consistency of the array distribution of bioactivity strength vs. the quasimolecular ion intensity of the modulating compounds on the paired chromatographic-based microassay arrays is established. A schematic diagram of the workflow used for the strategy is shown (Figure 2).

**Figure 2.**
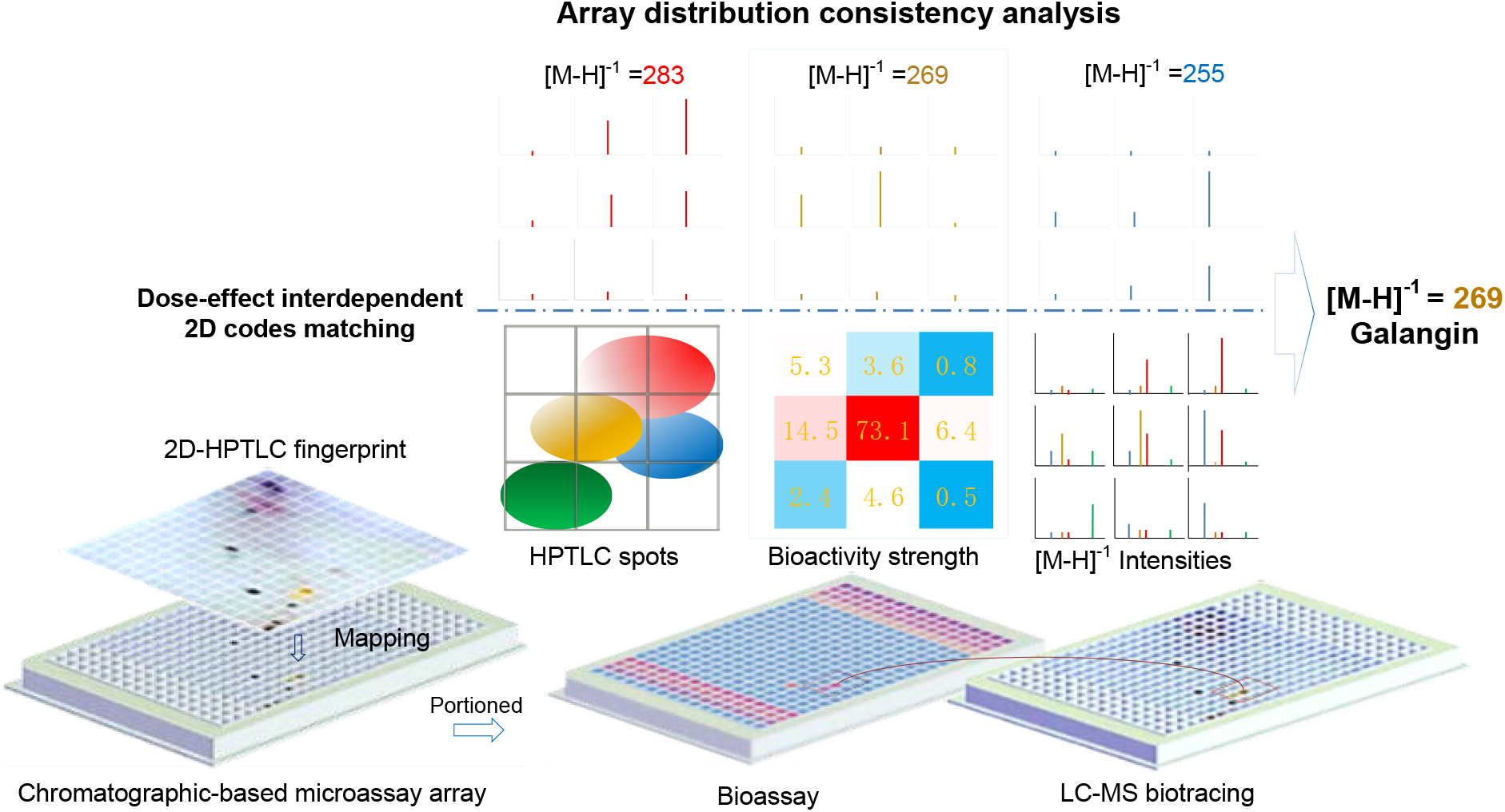
Schematic diagram of the workflow for establishing the array local self-consistency

In general, 2D-HPTLC may not be able to achieve complete separation of all the chromatographic spots of diverse compounds, and due to the shape, size and array distribution of HPTLC spots are irregular, each chromatographic spot does not always fall into a single microarray unit, and several chemical components may coexist in the same microarray unit. However, on the paired chromatographic-based microassay arrays, the chromatographic spots, even if they are partially overlapped, are differentiated by the microarray with vector feature. By virtue of the high resolution and high sensitivity of LC-ESI-MS, each digital specific quasi-molecular ion of coexisting small molecule compounds is characterized as its intensity array distribution gradient, and the trend difference of the array distribution gradients between the bioactivity strength vs. the digital-specific quasimolecular ion intensities of several coexisting components can be clearly distinguished in the corresponding regions (Figure 1). The bioactive compound must be present in those array units that exhibit bioactivity. When multiple components coexist there, it is important how to identify the molecule attributed to bioactivity from them. This experimental design provides a satisfactory solution.

Dose-effect interdependence is a basic pharmacological principle. Under this experimental design, the array distribution gradient between the bioactivity strength and the specific quasimolecular ion intensities of the respective modulating compound are local self-consistent, the attributes derived from the same compound will be auto-collimated, regardless of the irregular shape, size and array distribution of its chromatographic spot. Therefore, the experimental data obtained from the paired chromatographic-based microassay arrays are correspondingly extracted as homologous 2D code pairs, and are substituted into the array matching algorithm to determine whether there is dose-effect interdependence, and to identify the digital-specific molecular characteristics of the compound attributed to the bioactivity among the coexisting components in the corresponding array region [34, 35].

As a result, the target deconvolution, an important aspect of current drug discovery [36], is simplified as an array matching algorithm based on the principle of dose-effect interdependence, just like addressing constraint satisfaction problems (CSPs).

### Generation of 2D codes from obtained experiment results

This method was applied to galangal (*Alpinia officinarum* Hance, a famous traditional Chinese medicine) as a paradigm. The anti-cancer activity of galangal or its mechanism of action has been reported one after another [37, 38]. In the light of the anticancer effects of galangal, the bioassays of this experiment include the A549 and HepG2 cell viability assays and G-quadruplex ligand bioassay, and were performed on the chromatographic-based microassay arrays. The obtained bioactivity strengths are expressed in the array distribution and are then converted into two-dimensional codes by Excel (2010 version). The bioassay data and the generated two-dimensional codes are shown (**a, b** and **c** in Figure 3).

**Figure 3.**
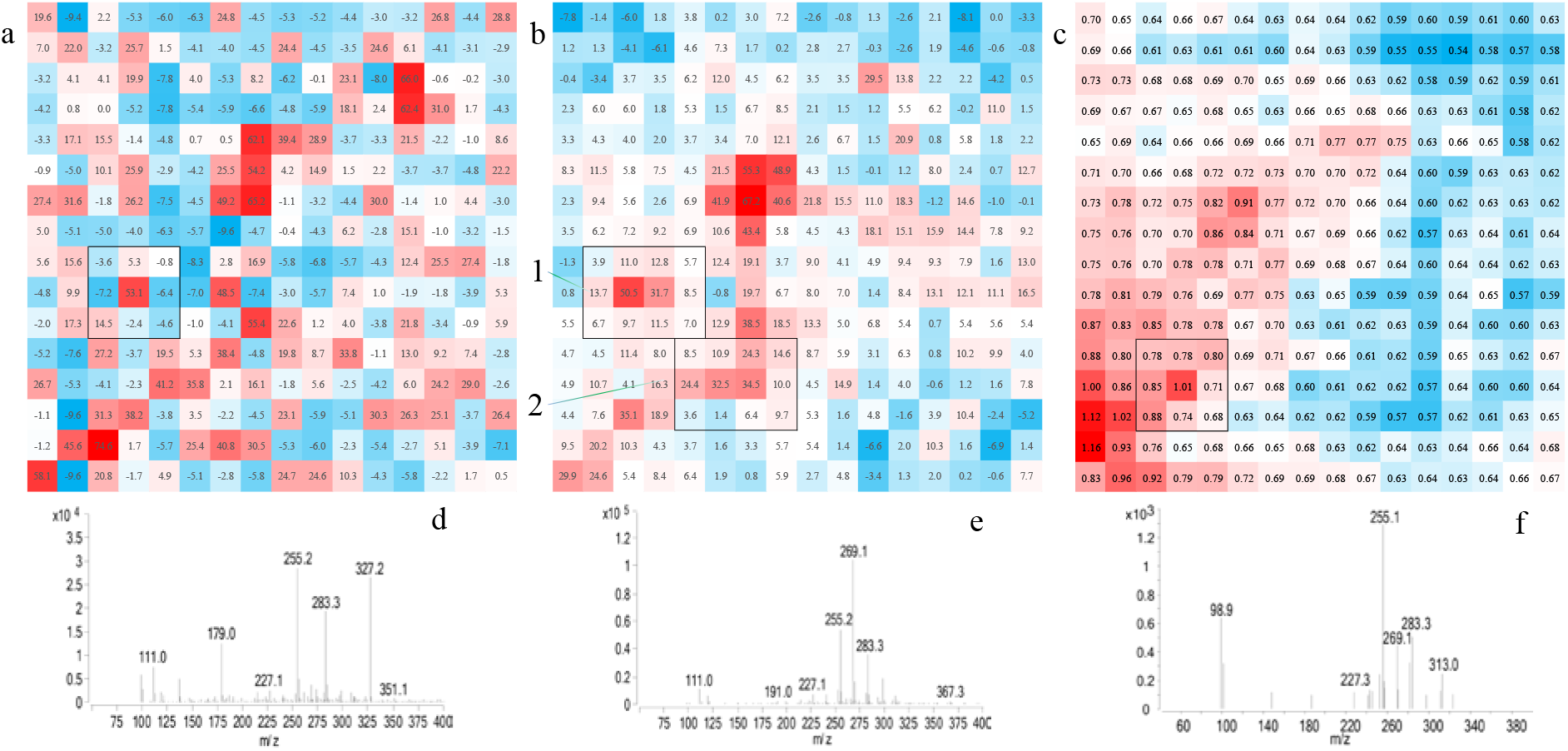
Array distribution of bioactivity strengths (2D code) generated from the results of the bioassay and the MS spectrum of the central array unit corresponding to the significant bioactive regions marked with a wireframe. (**a**) 2D codes generated from the viability data of the A549 cell line; (**b**) 2D codes generated from the HepG2 cell line; and (**c**) 2D codes generated from the G-quadruplex ligand bioassay. The MS spectra corresponding to the center array units of the significant bioactive regions marked with a wireframe in **a, b**2 and **c** are shown in **d, e, f**, respectively, and the MS spectra corresponding to **b**1 is similar to **d**.

Correspondingly, the compositions of the array units with significant bioactivity are further analyzed *in situ* by LC-MS and are highly resolved into their digital-specific quasimolecular ion intensities in the array distribution, and the MS spectra of the central array unit corresponding to the significant bioactive regions are obtained (**d, e**, and **f** in Figure 2, marked with wireframe. The chromatographic-based microassay arrays for the bioassay and LC-MS bioactivity tracing should be sampled from the same chromatographic-based microassay array stock sample as the homologous test samples). The main coexisting quasimolecular ions are determined based on the mass spectrum corresponding to the central array unit of the significant bioactive region. The array distribution data of the quasimolecular ion intensities of the main coexisting quasimolecular ions are extracted from the CSV file of the mass spectra of the array units corresponding to the significant bioactive region, and the array distribution of the quasimolecular ion intensity of each coexisting component is obtained. Then, the array distribution of the specific quasimolecular ion intensities is converted into the corresponding two-dimensional code by Excel (2010 version).

In this experiment, 2D-HPTLC has strong chromatographic distinguishing power and can even be used to identify isomers; 2D-HPTLC combining with LC-MS is a powerful means of directly identifying small molecular compounds [17, 18]. In addition, bioactive-directed LC-MS tracing on paired chromatographic-based microassay arrays is very effective because it is focused on several bioactive array units rather than on the entire microarray, the workload is greatly reduced.

### Dose-effect interdependence assessment

Dose-effect interdependence assessment of the array distribution gradients between the bioactivity strength and the specific quasimolecular ion intensities is performed with the CORREL (Array1, Array2) function in Excel (2010 version). The bioactive strength data array from the significant bioactive region and the adjacent region is selected as the variable Array1, while the quasimolecular ion intensity data arrays from the coexisting quasimolecular ions in the corresponding region are selected as the variable Array2. These experimental data are substituted into the CORREL (Array1, Array2) function as two-dimensional code pairs to assess the dose-effect interdependence. This method can be regarded as a prototype of the algorithm for this constraint satisfaction problem (CSP), which is used to evaluate the correlation between molecules and bioactivities in diverse compounds.

The data of the quasimolecular ion intensities, bioactivity strengths and the correlation analysis results obtained for the coexisting components in the four bioactive regions (**a, b**1, **b**2 and **c** in Figure 3, marked with a wireframe) are shown (Figure 4). We can visually observe which coexisting compound has an array distribution gradient of specific quasi-molecular ion intensity that tends to be consistent with the array distribution gradient of bioactivity intensity in the corresponding array region (Figure 4). Using array correlation algorithm, such as Excel CORREL function, to calculate correlation coefficient can make the array difference objective and quantitatively comparable, so as to facilitate computer-aided analysis. The absolute value of the correlation coefficient is used to rank the compounds corresponding to the coexisting quasimolecular ions for bioactivity screening. The larger the value, the more significant the difference and the higher the credibility. More obviously, if the array unit with the maximum quasimolecular intensity deviated significantly from the array unit with the maximum bioactive strength in the corresponding region, the corresponding compound would not be considered the bioactive compound.

**Figure 4.**
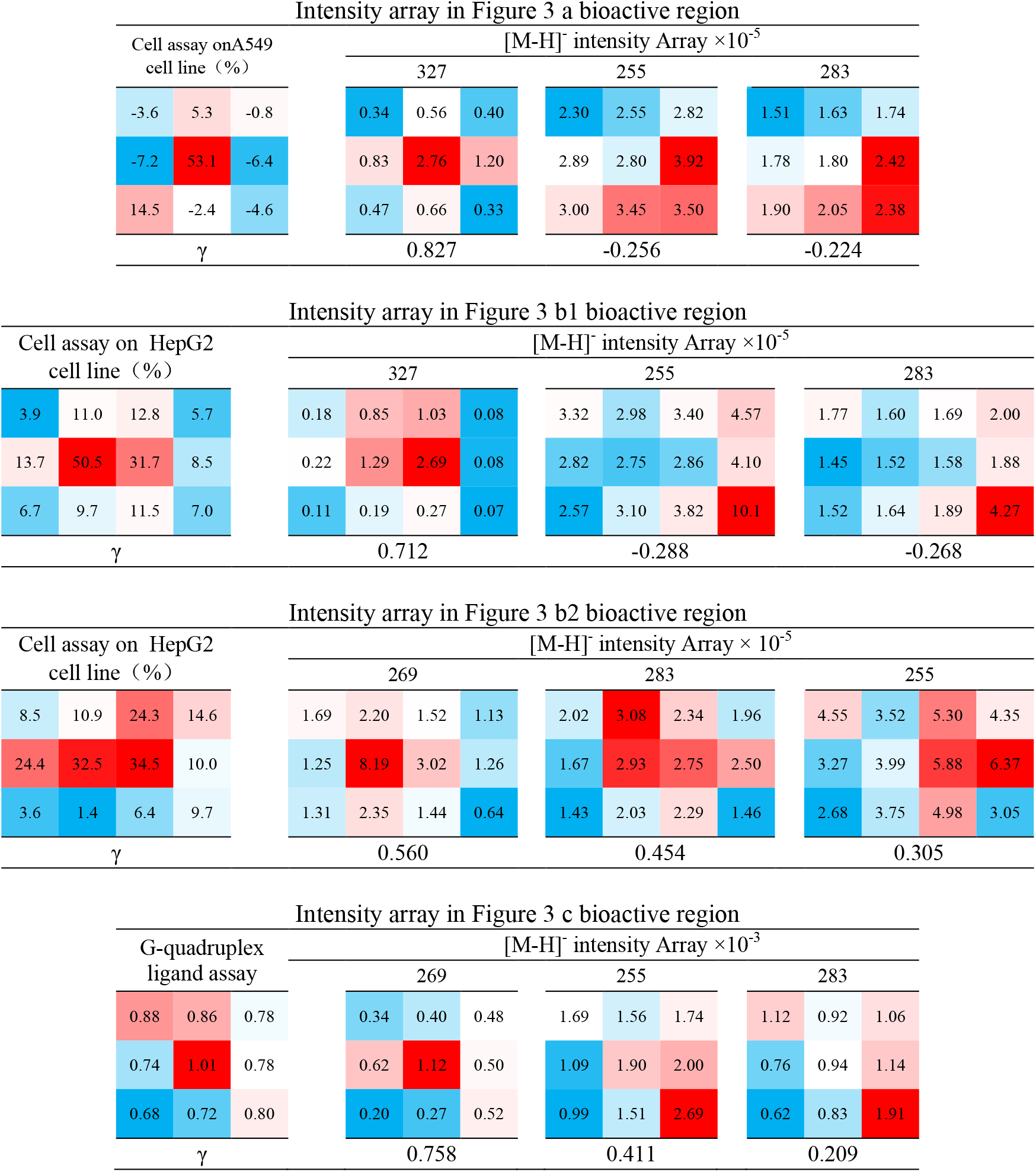
Dose-effect interdependence assessment of the array distribution gradients between bioactive strength and quasimolecular ion peak intensities. γ-Correlation coefficient of the array distribution of bioactive strength with the array distribution of quasi-molecular ion peak intensity obtained from the Excel CORREL function. The background color of the cells in the table is generated according to the cell values. The larger the value is, the darker the blue color; the smaller the value is, the darker the red color.

### Bioactive candidates and bioactivity evaluation

Based on the calculated correlation coefficients, compounds corresponding to 327 [M-H]^-^ and 269 [M-H]^-^ were identified as bioactive candidates by the cell viability assay, and 269 [M-H]^-^ was identified as a bioactive candidate by the G-quadruplex ligand bioassay. The bioactivity strength of the same quasimolecular ion in the array unit within the same bioactive array region was normalized to the contribution of this quasimolecular ion and incorporated into the bioactivity screening consideration. Since the small molecules causing bioactivity are identified from the original sample rather than from the reaction products, the abundant database of natural products can provide support. A literature search and a compound database search for bioactive candidates were performed. Referring to the literature [39], 327 [M-H]^-^ and 269 [M-H]^-^ in the galangal extract were speculated to belong to diphenylheptane A and galangin (CAS No. 68622-73-1 and 548-83-4), respectively.

The cell bioactivities of the two candidate compounds were evaluated by cell viability assays with 5-fluorouracil as a positive control. The cell growth inhibition curve data were analyzed with SPSS 18.0 to calculate the IC_50_ values. The IC_50_ values of diphenylheptane A (NICPBP, Lot No. 111757-200601), galangin (NICPBP, Lot No. 111699-200602) and 5-fluorouracil for the A549 cell line are 0.247, 0.089, and 0.023 mmol•L-1, respectively, and those for the HepG2 cell line are 0.259, 0.085, and 0.092 mmol•L-1, respectively. There was no significant difference between the bioactivity of galangin-induced G-quadruplex DNA formation (0.659±0.038, *n*=8) and that of aloe emodin (NICPBP, Lot No. 0795-9702), which was used as a positive control (0.686±0.026, *n*=8), at concentrations of approximately 0.5 μM.

### Resolution Analysis and Error Control

In order to reflect the array distribution gradient of the chromatographic spots in the 2D-HPTLC fingerprint by a chromatographic-based microassay array with a high degree of sharpness, the array unit is preferably designed to resemble chromatographic spots in size to avoid more than one separated chromatographic spots falling into the same array unit as a whole or the same chromatographic spot being split into similar array units in terms of chemical composition. It is feasible and practicable to characterize 2D-HPTLC fingerprint spots (typically approximately 2.5 mm to 4.5 mm in diameter) with commercial 384-square well high throughput microplates (with an array format of 4.5 mm×4.5 mm/well, 16 rows×24 columns), such as human cell-based phenotypic assays (typically 40 μL per well, with wells containing 1000 cells), target-based assays and LC-ESI-MS biotracing. The array distribution of the chromatographic spots (of the compounds developed with the sulfuric acid-vanillin reagent) in a 384-well array grid is shown (Figure 5).

**Figure 5.**
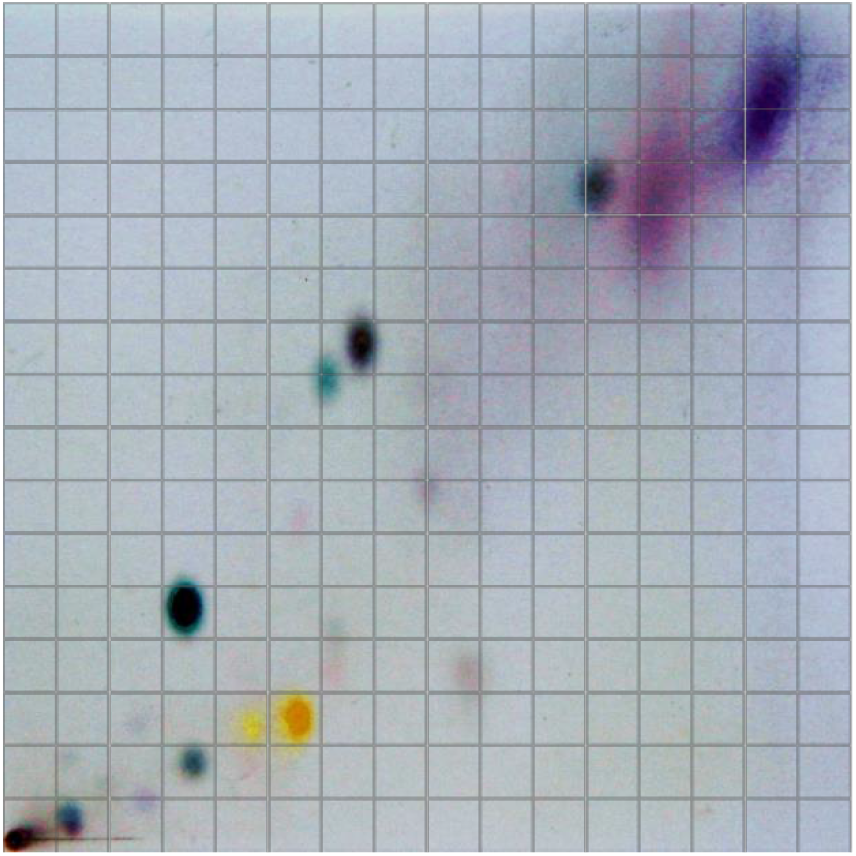
Schematic diagram of the array distribution of the 2D-HPTLC chromatogram of galangal extract

Cell viability assays are the bottleneck of the resolution, which needs a certain space for cell growth, and the components of chromatographic spots should reach the drug dose range. As the RSD of the Alamar Blue cell viability assay used in this experiment is about 8%, this requires that the cell viability of the detected array units is more than 24% (3 times the RSD). Since the content of chromatographic components in the array unit may fluctuate between 1 ∼ 1/4 (in the worst case, one spot may be evenly distributed in four array units), so the proportion of non-overlapping parts of chromatographic points must be greater than about 25%. This means that the resolution ratio of chromatographic spots should be greater than 0.5 (As the resolution ratio reaches 1.5, the adjacent two spots reach baseline separation). In terms of planar chromatography technology, such resolution requirements are perfectly achievable.

In fact, the worst case mentioned can be avoided. Since on the HPTLC plate (silica gel 60, GF254) used in this experiment, most of the chromatographic spots of biomolecules can be detected non-destructively under UV light, and the bioactive regions can be known during pre-bioassays for sample loading amount, so the bioactive regions can be pre-located in as few array units as possible by adjusting the relative position between the array grid and 2D-HPTLC fingerprint.

### The scientificity and effectiveness of the strategy

In this strategy, a pair of chromatographic-based microassay arrays are created to characterize and assemble the modulating bioactivity and molecular characteristics of the compounds separated on 2D-HPTLC into self-consistent array distribution gradient, the contribution of small molecules to bioactivity was assessed according to the basic pharmacological principle of dose-effect interdependence, and all the experiments employed the conventional proven pharmaceutical research methods and specifications. Ultimately, the screened candidates were determined by bioactivity assessment compared with bioactive control. Therefore, the results obtained by the strategy are scientific and valid.

Due to the combined multidisciplinary approaches, this strategy has achieved a lot of research innovations. For the first time, the total compounds separated on the HPTLC fingerprint were subjected to mammalian cell-based phenotypic assay without interference from the chromatographic matrix; the observation indicators are converted from conventional chromatographic spots to array distribution gradients of essential substance attributes, e.g. apoptotic cell morphology and digital-specific quasimolecular ion, and the resolution is greatly improved; The compounds’ attributes are characterized in intensity array distribution gradient and digitized into homologous 2D codes, so the small biomolecules can be recognized through array matching algorithm with the assistance of computer. Therefore, the separation and *in situ* recording of chemical constituents on TLC, the labeling of specific bioactivities and the high resolution of mass spectrometry are combined for a parallel and synergistic comprehensive screening, which greatly reduces interference and significantly improves the resolution. This avoids the situation that diverse compounds of a complex samples have to be separated to monomers regardless of whether they are bioactive, and avoids the loss of potential bioactive compounds in tedious fraction cutting and concentration preparation without bioactivity monitoring. In this way, high-throughput digital preliminary screening of small biomolecular in diverse compounds can be realized, and the efficiency is significantly improved and the workload is greatly reduced.

Another notable aspect, this experiment shown that galangin in *Alpinia officinarum* Hance has bioactivities for inducing G-quadruplex DNA formation and inhibiting cancer cell proliferation, while diphenylheptane A also inhibits cancer bioactivity. These results suggest that galangin induces the formation of G-quadruplex DNA and is involved in the inhibition of cancer cell proliferation, which may be one of the pathways of galangin’s anticancer activity. Although diphenylheptane A also has anticancer activity, its mechanism of action is different from that of galangin.

It is important to “integrate both the phenotypic and target-based approaches to estimate a relevant network from compound to phenotype in screening” [7, 8]. In this experiment, human cell phenotypic (two cell lines) and ligand based bioassay were coupled with LC-ESI-MS biotracing on the paired chromatographic-based microassay arrays. By considering the digital-specific quasimolecular ions as nodes and dose-effect interdependent code pairs as links, the cell phenotypic and target-based bioassays can be integrated with LC-ESI-MS biotracing to establish an extendable local self-consistent framework. Thus, the cell-based phenotypic assay, target-based assay and small biomolecules can be comprehensively correlated. This helps to eliminate false positives in screening and understand the mechanism of action, thereby significantly improving the reliability of small biomolecule identification.

### Factors affecting self-consistent analysis

In this experiment, the correlation coefficient is not as large as the normal standard calibration curve, and some microarray regions that appear to exhibit bioactivity fail to resolve molecules attributed to bioactivity (Figure 3). There may be particular reasons, such as the high content, the linear range of the signal response, and the memory effect of the thin-layer silica gel. The components of natural products are diverse and vary greatly in content. The concentration levels of TLC spots prepared using a certain sampling amount are not all suitable for bioassays, and the ultrahigh concentration levels in some microarray units may lead to nonspecific bioactivity effects. In addition, these phenomena can occur if the local content is too high due to spot overlap. The correlation coefficient here reflects the multifactorial correlation between the response signals of the bioassay system and those of the LC-ESI-MS system. If one of the respective signal responses detected by the bioassay or LC-ESI-MS is not within the linear response range, the correlation may weaken. In addition, due to the memory effect, trace components remaining in the thin-layer silica gel can be strongly eluted by methanol. The interferences from these components will appear as noise detected by LC-MS or bioassays. This type of memory residue of the complex components is not the same, and interference cannot be completely deduced from the reagent blank. After all, the chemical composition on the array unit of the chromatographic-based microassay array is usually not a single component, and it also involves multi-factor response values. The correlation coefficient mentioned here is different from the correlation coefficient of the standard calibration curve obtained for the standard reference material.

## Conclusions

Through building the paired chromatographic-based microassay arrays, high-throughput micro-separation, *in situ* recording, μTAS, coding interaction and correlating recognition of substances are achieved, and an extendable local self-consistent framework that combining mammalian cell phenotypic and target-based bioassay with HPTLC-MS is established. Therefore, the small molecular contributions and correlations of bioactivities and their pathways can be associated for comprehensive screening. This approach can make use of abundant resources, such as validated drug research methods, drug databases and TLC fingerprint library recorded in pharmacopoeia. The digital acquisition and recognition algorithm of chromatographic data is conducive to the computer-aided processing of big data, and has the prospect of artificial intelligence. This research strategy will greatly promote the drug discovery of small molecules.

## Supplementary information

The information includes preparation of the metal array grid interface, preparation of the square chip array of the silica gel layer, and elution of the silica gel chips with a 384-square-well filter plate. A schematic diagram of preparation of chromatographic-based microassay array can be found in the online Supporting Information. Figures S1−S3 are shown.

## Author contributions

Yifeng Deng conceived the project, designed all the experiments in this study and drafted this manuscript for publication. In addition, he performed all the experiments for the G-quadruplex ligand bioassay and consistency analysis of all the experimental data. Zhenpeng Lin performed the cell viability assays and the LC-MS analysis, processed the related data, prepared some Chromatographic-based microassay arrays, and created some of the CAD drawings. Yuan Cheng carried out the preliminary experiments for this study.

## Acknowledgments

We thank Yu Zhong (Analysis Centre of Guangdong Medical University, China) for technical assistance during LC-MS analysis, Liubo Lan for help with the initial cell viability assay, and Ning Li (Affiliated Hospital of Guangdong Medical University, China) for providing the A549 and HepG2 cells.

## Notes

The authors declare that they have no competing financial interests.

## Supporting Information

### S1 Preparation of the metal array grid interface

A design drawing of the metal array grid interface is shown (Figure S1). The metal array grid is made of 3D printed Co-Cr alloy (OEM according to drawings). The grid array format is the same as that of the 384-well microplate, with a format of 4.5 mm×4.5 mm square and horizontal 24×16 vertical arrays. The grid space is a regular inverted quadrangular pyramid. The vertical section of the grid frame is an isosceles triangle with a height of 2.5 mm. The width of the base of the isosceles triangle is the same as the wall thickness (approximately 0.70 mm) of the well of the 384-square-well microplate.

**Figure S1.**
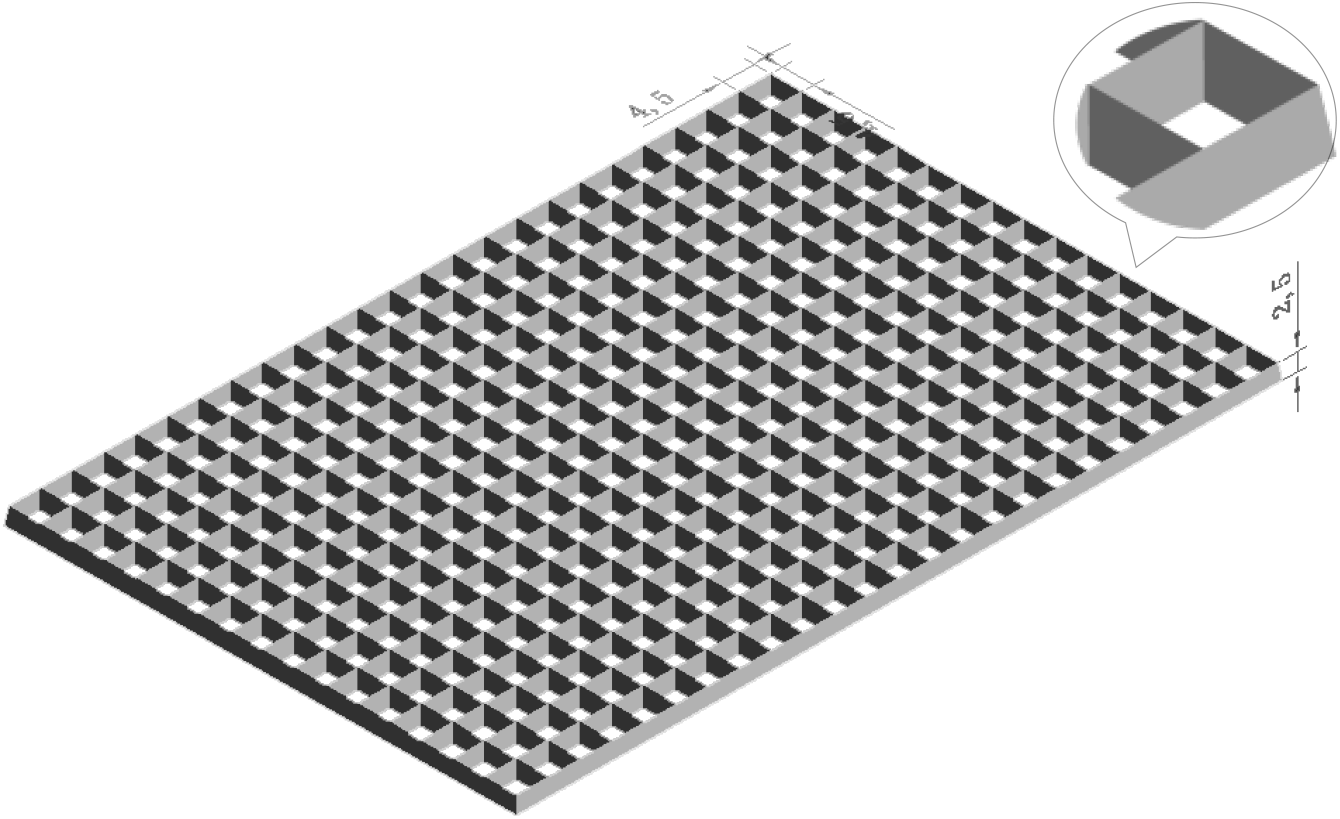
Metal array grid interface design

### S2 Preparation of the square chip array of the silica gel layer

The silica gel layer of the chromatographic region of a sample’s 2D-HPTLC fingerprint was split by the metal grid interface (S1) at a parallel pressure and stripped from a thin aluminum sheet into a flat square chip array. Specifically, the components of a sandwich composed of a silicone rubber sheet (thickness 2 mm), metal grid interface, PVDF membrane (Millipore, thickness 0.2 mm), dried HPTLC aluminum sheet and paper (PVDF film protection paper) were laminated in order from bottom to top. The sharp edges of the metal grid interface were aligned and faced the thin layer of silica gel across the PVDF membrane; then, the edges were pressed with a parallel pressure of about 8×10^3^ kPa (Specac Atlas® Manual Hydraulic Press-15T). When the thin ductile aluminum sheet and the thin crisp layer of silica gel touched the sharp square opening edge of the grid interface with pressure, the thin layer of silica gel was split into square chips and was stripped from the thin aluminum sheet, thereby adhering to the PVDF membrane, which was slightly embedded in the metal grid interface. The stripping rate of the silica gel thin layer reaches 91.3%, and the chips of the silica gel thin layer are arrayed regularly and orderly as a square chip array.

### S3 Preparation of the paired chromatographic-based microassay array

The openings of the 384-square-well filter plate (Pall 5072) were aligned with the array of silica gel layer chips adhered to the PVDF membrane (slightly embedded in the metal grid interface), and the PVDF membrane was touched with a 16-channel pipette (without the pipette tip) from the other side of the metal grid interface so that the silica gel layer chips fell into the corresponding wells of the 384-square-well filter plate (Figure S2). The components in the silica gel chips were eluted with methanol into another microplate. The microplate containing the eluent was evaporated to dryness on a vacuum centrifuge concentrator (269 ×g, 50°C, -80 kPa, 1.5 hours) and prepared as the stock chromatographic-based microassay array. The paired chromatographic-based microassay array with the same chromatographic distribution is prepared by transferring a portion of the sample from the stock chromatographic-based microassay array to the corresponding array unit of another 384-well microplate.

**Figure S2.**
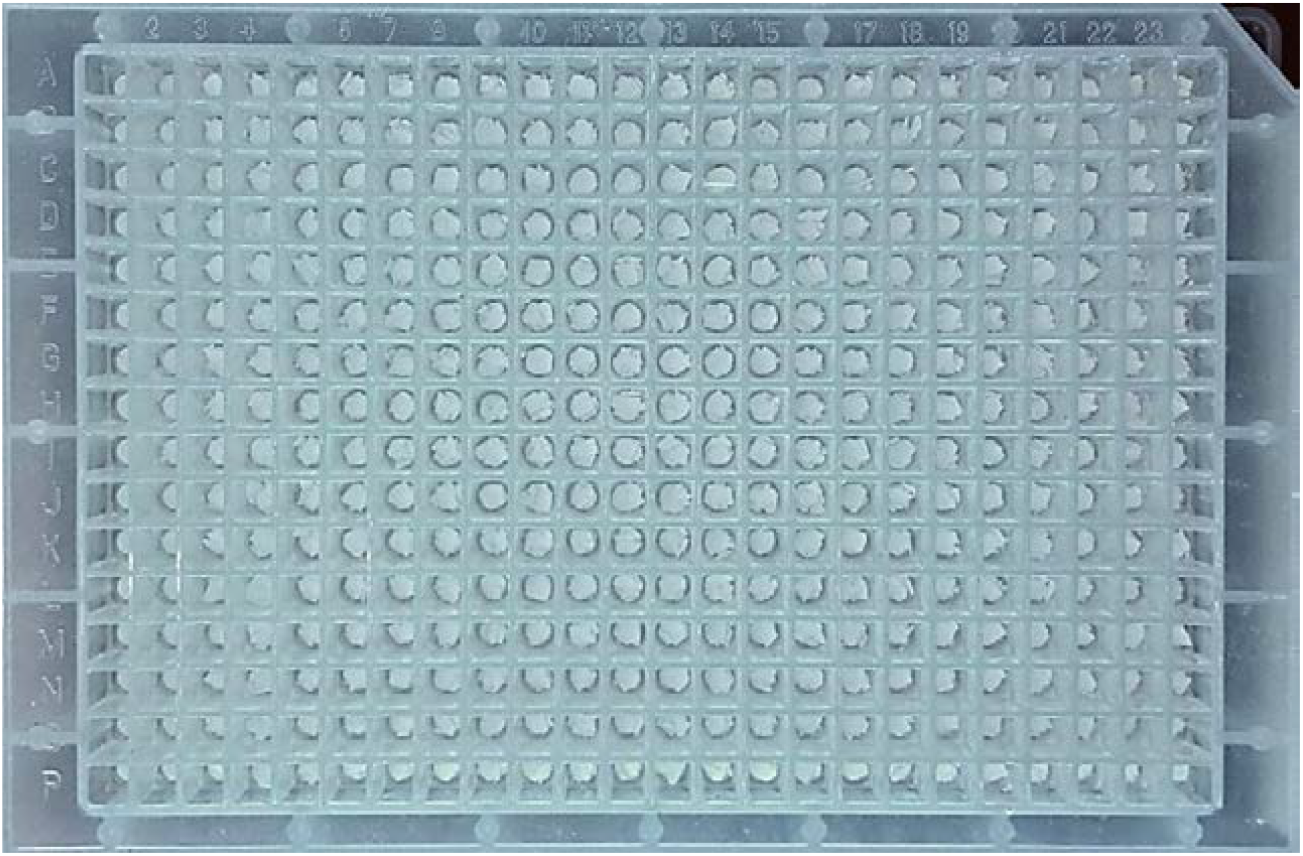
The components in the silica gel chips were eluted with a 384-square-well filter plate. The square chips of the silica gel layer were positioned into the corresponding 384-square-well filter plate

### S4 Schematic diagram of preparation of chromatographic-based microassay array

A schematic diagram of preparation of Chromatographic-based microassay array is presented (Figure S3).

**Figure S3.**
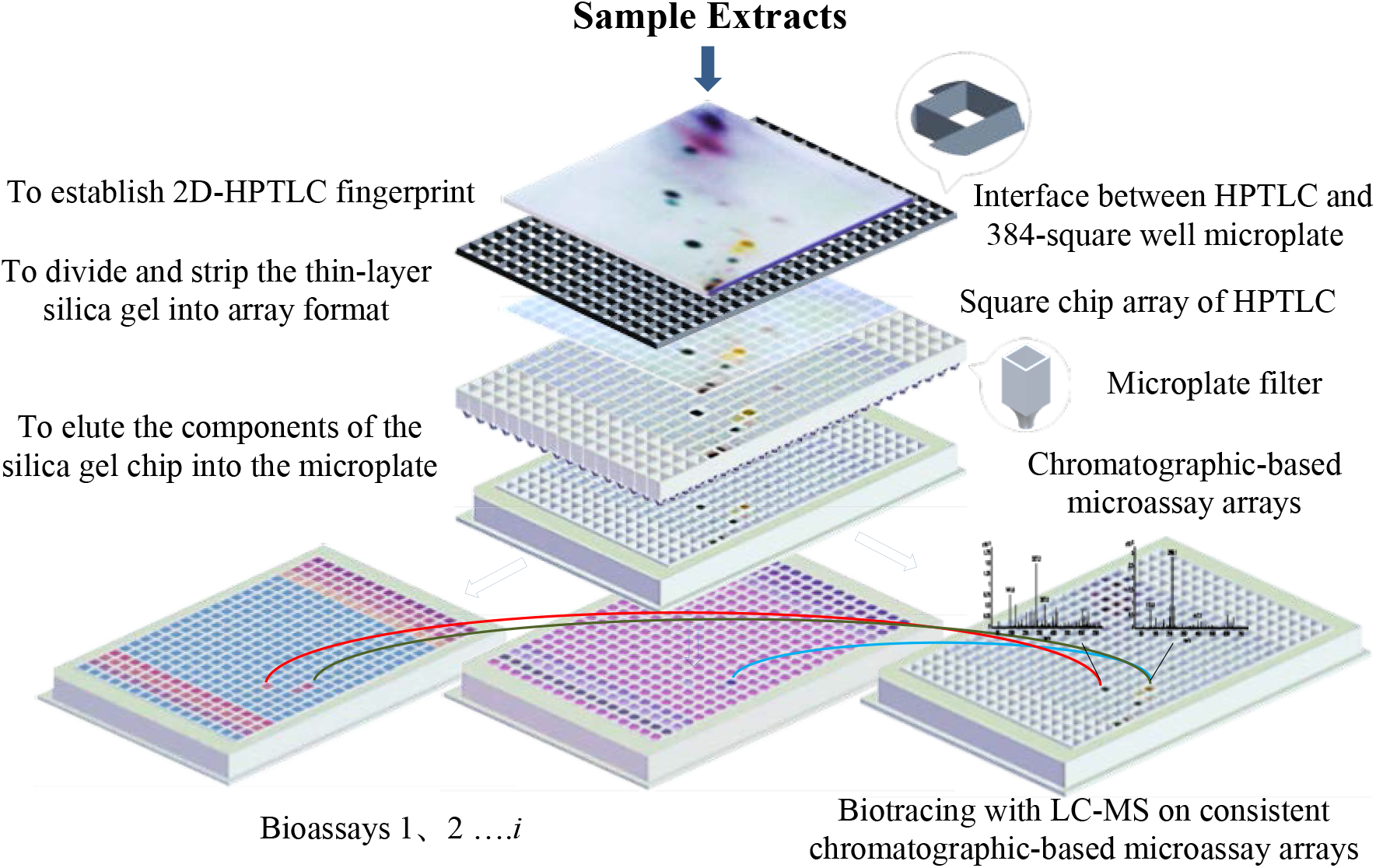
Schematic diagram of preparation of chromatographic-based microassay arrays

